# Analyzing multisensory integration: do’s and dont’s

**DOI:** 10.64898/2026.04.15.718785

**Authors:** Haocheng Zhu, Ulrik Beierholm, Ladan Shams

**Affiliations:** Department of Psychology, University of California, Los Angeles, USA; Department of Psychology, University of Durham, UK; Department of Bioengineering, University of California, Los Angeles, USA; Interdepartmental Neuroscience Program, University of California, Los Angeles, USA

**Keywords:** multisensory integration, Bayesian causal inference, perceptual inference, crossmodal illusions, computational modeling

## Abstract

Multisensory perception is a cornerstone paradigm for understanding how the brain constructs coherent representations of the world from noisy, fragmented sensory inputs. For decades, researchers have used the magnitude of crossmodal illusions, the width of the temporal binding window, and related behavioral indices as direct proxies for ’integration strength,’ and have leveraged these measures to compare multisensory function across developmental, clinical, and aging populations. Here we argue that this descriptive practice is fundamentally compromised: behavioral readouts of multisensory integration are composite measures jointly shaped by unisensory precision, amodal priors, and the binding process itself, and cannot be interpreted in isolation. Drawing on simulations within a Bayesian Causal Inference framework, we show how identical behavioral patterns can arise from very different underlying causes, leading to systematic misattribution of group differences to ’deficits’ or ’enhancements’ in integration. We review complementary computational frameworks, including drift diffusion, multisensory correlation detection, and statistical facilitation models, and outline their respective explanatory limits. Finally, we provide a model-based inference pipeline, from experimental design and unisensory baselines to parameter estimation and interpretation, that disentangles sensory fidelity, prior expectations, and integrative tendency. Adopting this normative approach is essential for cumulative progress in basic multisensory research and for its translation to neuropsychiatric assessment, lifespan research, and artificial perceptual systems.

## 1 Introduction

A fundamental challenge for any intelligent system, biological or artificial, is to construct a coherent representation of the world from noisy, fragmented sensory inputs. Multisensory perception (MSP) serves as a paradigmatic testbed for understanding this process, requiring the nervous system to solve a complex problem of probabilistic inference: evaluating the causal probability that signals belong together, and dynamically weighting inputs based on their reliability.

For decades, this field has been anchored by classic behavioral paradigms, from the McGurk effect in speech perception (McGurk & MacDonald, 1976) and the ventriloquist effect in spatial localization (Bertelson & Aschersleben, 2003; Alais & Burr, 2004), to the rubber hand illusion (Botvinick & Cohen, 1998; Ehrsson et al., 2004) and the sound-induced flash illusion (Shams et al., 2000, 2002). These behavioral paradigms are widely employed to probe distinct aspects of multisensory processing. That led many researchers to use such paradigms and their respective measures of MSP as a basis for comparing one’s integration abilities across different populations and conditions. From that, an important line of such research has been oriented toward dimensions of comparison, for instance, the integration processes between typically developed persons and those with autism spectrum disorders. Indeed, such studies have revealed statistically significant differences in multisensory processing in autism, with findings for example, showing decreased audiovisual temporal binding window and distinct multisensory temporal processing (Wallace & Stevenson, 2014; Stevenson et al., 2014; Noel et al., 2018). Other dimensions of MSP have similarly been compared across clinical populations, such as in individuals with schizophrenia, where altered temporal integration windows compared with healthy controls were reported (Williams et al., 2010; Zhou et al., 2022). Another important axis of comparison concerns developmental trajectories. From childhood through adulthood and into older age, both the extent and the strategy of cue combination can change. Younger children typically exhibit broader audiovisual temporal binding windows and a different weighting of modalities relative to adults, whereas older adults often show patterns consistent with reduced temporal precision and altered reliability-based weighting (Nardini et al., 2008; Gori et al., 2008; Chen et al., 2016; Chan et al., 2014). Experience-dependent factors, including training, environmental exposure, and domain-specific expertise, can also reshape MSI, demonstrating measurable plasticity across the lifespan (Shams & Seitz, 2008; Stevenson et al., 2012).

However, the utility of MSP as a model system for perceptual inference is currently compromised by the reliance on descriptive heuristics. A common methodological approach has been to infer ‘stronger’ or ‘weaker’ integration from the magnitude of multisensory illusions or the width of the so-called temporal window of integration (TWI) (e.g., Stevenson et al., 2012; Zhou et al., 2018). This inference rests upon the often-untested premise that the degree of binding is the only, or principal, determinant of these behavioral outcomes. Indeed, recent reviews have highlighted conceptual ambiguities in how the TWI has been defined and operationalized, noting that measures based on synchrony judgments may reflect a distinct construct (Jertberg et al., 2025). Behavioral indices are fundamentally composite measures, shown to be confounded by factors not specific to the core integrative mechanism, including the sensory precision of unisensory inputs and amodal priors (Odegaard & Shams, 2016; Zhu et al., 2024; Dong et al., 2025; Beierholm et al., 2025). For instance, a "stronger" illusion in one group may simply reflect poor visual acuity rather than a genuinely larger probability of integrating the audiovisual stimulus (Zhu et al., 2024). Likewise, a wide TWI in older adults or certain clinical conditions is often attributable to one modality’s impaired temporal resolution rather than an increased integrative tendency (Odegaard & Shams, 2016).

Particularly, differences in the specific subtypes of the sound-induced flash illusion, such as ‘fission’ (perceiving more events than presented) and ‘fusion’ (perceiving fewer events) effects in the task of temporal numerosity judgment, have been interpreted as evidence for separate multisensory mechanisms. This interpretation has been challenged with novel computational frameworks, especially those rooted in Bayesian Causal Inference (BCI), showing that both classes of illusions result from one integrating process driven by unisensory reliability and prior expectations, not necessarily involving different integrating circuitry (Körding et al., 2007; Shams & Beierholm, 2010; Fetsch, DeAngelis, & Angelaki, 2013; Rohe & Noppeney, 2015; Odegaard, Wozny, & Shams, 2015; Zhu et al., 2024; Dong et al., 2025). Under BCI, changes in illusion magnitude or TWI cannot be attributed to multisensory processes unless they are first separated from unisensory precision and amodal priors. This disentangling is essential for group comparisons (ASD vs. typically developing, healthy vs. schizophrenia, and across age cohorts; Rohe et al., 2024). Without such disentangling of unisensory processing, amodal expectations, and multisensory binding process by meticulous modeling and baseline measures, the observed differences may be uncritically considered multisensory integration ‘deficits’ or ‘enhancements’ when actually reflecting changes in the unisensory acuity or broader perceptual strategies. To unlock the full explanatory power of multisensory paradigms, the field must therefore move beyond these descriptive heuristics to a normative computational framework that explicitly disentangles sensory fidelity from integrative priors.

In the following sections, we will discuss common pitfalls in interpreting MSP measures, highlight how unisensory and amodal factors can confound conclusions, and propose strategies, ranging from computational modeling to comprehensive experimental designs to more rigorously assess and interpret results in both research and applied contexts. Building on these claims, we provide a model-based inference pipeline from design to analysis and a minimal, reproducible model specification that directly informs task planning, parameter estimation, and interpretation.

## 2 Common Pitfalls in Interpreting Behavioral Indices of Multisensory Integration

Behavioral paradigms used to quantify multisensory integration typically rely on multisensory illusions, cross-modal biases, and temporal judgments. These indices have substantially advanced our understanding of how sensory signals are integrated; however, when interpreted in isolation they are prone to misinterpretation. It is common to treat illusion magnitude, the breadth of the temporal binding window, or the presence of distinct illusory outcomes as direct readouts of the strength or nature of integration. Yet a growing theoretical and empirical literature shows that these measures are also shaped by factors other than the integrative mechanism itself, including unisensory precision and amodal priors.

Three such pitfalls regarding interpreting behavioral data associated with MSP are summarized below. They are oft-repeated, but these make a specific case for a broader, more rigorous analytical approach concerning unisensory reliability, learned expectations, and the processes of probabilistic inference that shape observed outcomes.

### 2.1 Illusion Magnitude Does Not Necessarily Index Integrative Strength

One common assumption is that the magnitude of a multisensory illusion, such as the degree of auditory-driven visual misperception, is tantamount to the strength of the binding between the senses. Several studies have conceptualized classic multisensory illusions, such as the McGurk effect, sound-induced flash illusion, or ventriloquist effect, on such grounds, often concluding that increased susceptibility to such illusions refers to "enhanced" multisensory integration (e.g., de Boer-Schellekens & Vroomen, 2014; DeLoss, Pierce, & Andersen, 2013; Stevenson et al., 2012; Tiippana, 2014).

However, while large illusory magnitudes could potentially stem from more robust integration, this may not be necessarily the case. Indeed, the measured magnitude of an illusion is subject to significant modulation through the following: (1) Unisensory Reliability: Reduced precision in one modality may make it more easily swayed by another, thus generating a larger illusion at no real increase in integrative propensity (Figure 1A illustrates how a large versus small difference in unisensory reliabilities shifts the auditory posterior toward the visual estimate, even when the integration mechanism itself is held constant); (2) Amodal Priors: An observer’s prior bias for a specific value of source (to be estimated) such as number of flashes (in case of the sound-induced flash illusion, or a spatial location (in case of the ventriloquist illusion) may inflate the illusion’s magnitude. (3) Binding Tendency: Even when unisensory precisions are held constant, a higher prior probability that the two signals share a common cause pulls the posterior estimate more strongly toward a fused percept, amplifying the illusion independently of both sensory noise and amodal priors (Figure 1B). A stronger illusion, therefore, does not necessarily constitute an actual change in the integrative process, as any of these three factors,alone or in combination, can modulate the observed illusion magnitude.

**Figure 1.**
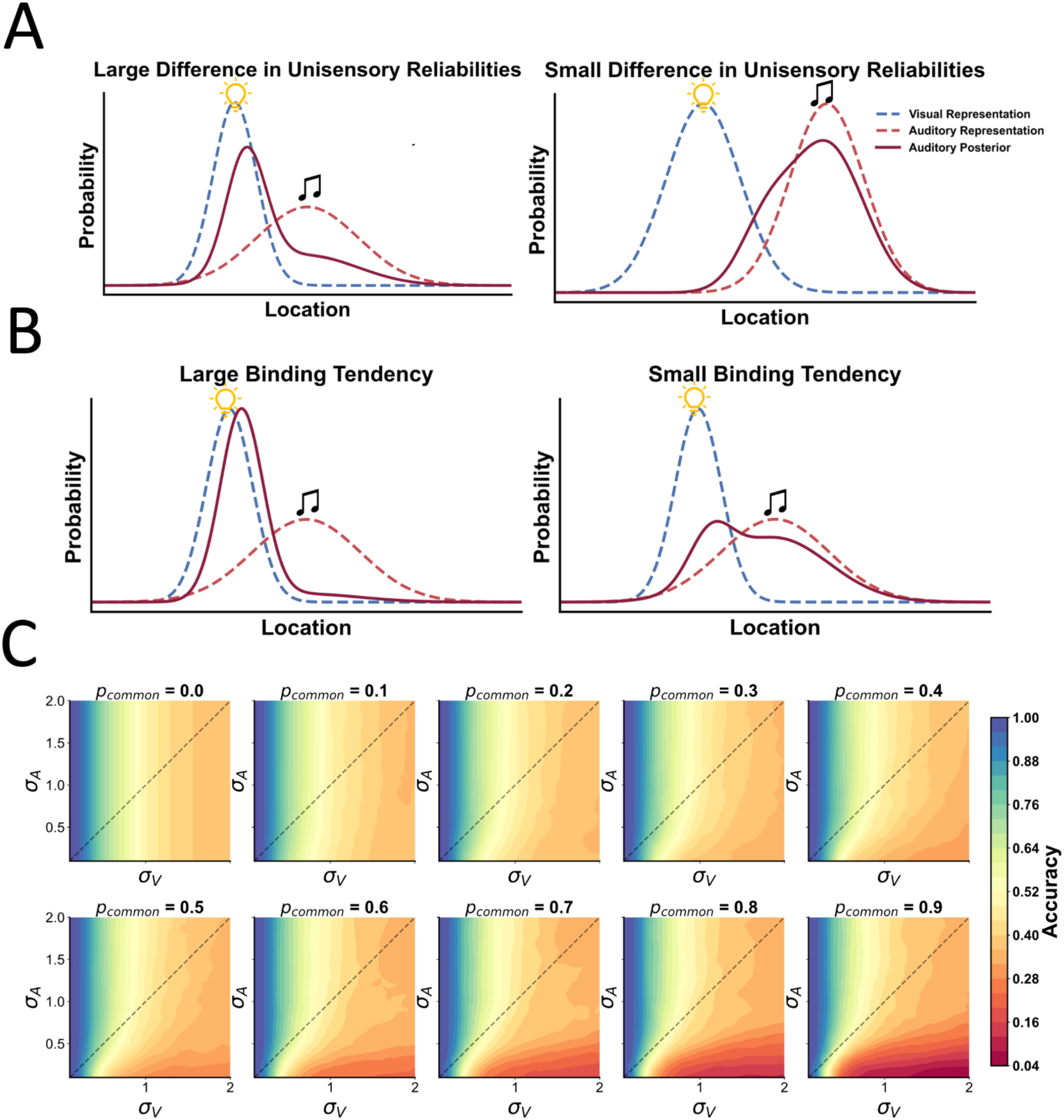
Disentangling unisensory reliability and binding prior in BCI. **(A) Simulations showing the role of relative unisensory reliabilities, and integration tendency in observed crossmodal bias** Panels illustrate how relative unisensory reliabilities and the binding tendency (*p_common_*) shape cross-modal bias under a Bayesian Causal Inference regime in an auditory spatial localization task. Simulations were generated using BCI Python Toolbox (Zhu et al., 2024). The x-axis denotes stimulus location; the y-axis denotes probability. Dashed curves are the visual (green) and auditory (purple) sensory likelihoods; the solid orange curve is the auditory posterior. Top row: Holding the binding/integration tendency fixed, a large reliability mismatch (vision ≫ audition; left) causes the posterior to shift strongly toward vision (i.e., a large bias); whereas similar reliabilities (right), leads to a small shift (i.e., small bias). (B) Holding reliabilities fixed, a higher *p_common_* yields a larger shift (left), whereas a lower Pcommon results in a small bias(right). These schematics demonstrate how each of these factors (unisensory reliabilities, and binding tendency) can have analogous effects on observed crossmodal bias, also known as the Ventriloquist illusion. **(C) Predicted accuracy under model averaging for the F1B2 condition of the sound-induced flash illusion as a joint function of audiovisual reliabilities and causal prior.** Panels show Bayesian Causal Inference simulations in which the probability of a common cause (*p_common_*) is swept from 0.0 to 0.9 (left-to-right, top-to-bottom). Within each panel, the x-axis denotes the visual noise (*σ_V_*) and the y-axis the auditory noise (*σ_A_*). Contours are coloured according to the model-predicted proportion of correct judgements (see scale bar). Decreasing *P*common or decreasing either sensory variance systematically elevates accuracy, while asymmetric changes in *σ_V_* versus *σ_A_* produce characteristic tilts in the iso-accuracy bands, reflecting the model’s reliability-weighted integration of the two modalities.

Several studies on audiovisual speech show this problem. For example, Lee et al. (2024) investigated the McGurk effect, whereby an auditory ‘ba’ paired with visually presented ‘ga’ results in subjects perceiving either ‘da’ or ‘tha’. They concluded that the overall fusion rate is impervious to extensive music training and therefore reflects a stable, population-level index of audiovisual integration. While this is plausible to a degree, group differences in visual clarity or in the relative weight observers place on prior phonetic expectations can just as easily drive variation in McGurk responses, without necessarily implicating changes in the underlying integration mechanism. More recently, investigators have begun to isolate these independent factors; for example, Dong et al. (2025) systematically examined how manipulating unisensory variance alone modulates susceptibility to the McGurk illusion.

In the sound-induced flash illusion, Shams et al. (2000, 2002) first demonstrated that multiple beeps accompanying a single flash could lead observers to report perceiving multiple flashes. While the original studies addressed the unisensory and integrative factors critically, several follow-up studies have since adopted the ‘number of illusory flashes’ as almost the exclusive measure of multisensory integration strength (e.g., Setti et al., 2011; Kheirkhah et al., 2022; O’Dowd et al., 2023; Hirst et al., 2019; Sun et al., 2022). O’Dowd et al. (2023) reported that older adults whose grip-strength trajectories placed them in the ‘low-strength’ cluster were significantly more susceptible to the sound-induced flash illusion (SIFI) at longer audiovisual stimulus-onset asynchronies (SOAs; 150 and 230 ms) than higher-strength peers, consistent with a broadened temporal binding window (TBW). However, because the study did not separately estimate unisensory temporal precision, amodal priors, or decision strategies (Wozny et al., 2010), the elevated illusion rate cannot be uniquely attributed to altered integration; it may equally reflect noisier unisensory timing or differences in numerosity priors. In this respect, such an interpretation overlooks the possibility that noisier visual representations (i.e., reduced visual reliability) make observers more susceptible to auditory capture of vision, thereby amplifying the illusory percept even if the underlying multisensory computations remain unchanged. Similarly, in the Ventriloquist paradigm, an increased shift of perceived visual location toward an auditory cue has sometimes been interpreted as signifying ‘greater’ audiovisual integration (eg. Bertelson & Aschersleben, 2003; Lavan et al., 2022; Park et al., 2021). However, follow-up modeling work has suggested that variable unisensory precision, especially the visual modality, can account for a large part of the variance observed in the ventriloquist effect (Körding et al., 2007; Wozny et al., 2010; Rohe & Noppeney, 2015). Individuals with less precise visual localization tend to show a larger ventriloquism bias even without any genuine increase in the propensity to integrate cues. Attributing a higher illusion magnitude to ‘stronger integration’ therefore risks conflating integrative mechanisms with unisensory or amodal factors. A more rigorous approach is to measure unisensory precision and fit models that include priors, thereby isolating the multisensory component from variability in sensory reliability and perceptual expectations (e.g., Shams & Beierholm, 2022; Rohe & Noppeney, 2015; Rohe et al., 2019).The unisensory baseline data and the application of computational frameworks, such as Bayesian Causal Inference, help researchers parse out to what extent illusion magnitude is reflective of true crossmodal binding rather than other sources of perceptual variability.

To further illustrate how both unisensory factors and multisensory priors jointly shape behavioral outcomes, we employed the BCI toolbox recently developed by Zhu et al. (2024). Specifically, we simulated the classic single-flash–double-beep configuration of the sound-induced flash illusion under ten discrete values of integration tendency (*p_common_* ranging from 0.0 to 1.0) alongside systematic variations in visual and auditory precisions (noise parameters *σ_V_* and *σ_A_*). As shown in Figure1C, the resulting simulations clearly demonstrate that changing either unisensory reliability (i.e., the *σ* parameters) or the integration tendency (*p_common_*) can substantially alter the prevalence and strength of the illusion. Thus, even when integrative mechanisms (captured by *p_common_*) remain fixed, unisensory precision alone can amplify or diminish illusory perceptions, and conversely, high *p_common_* can yield strong illusions even under relatively modest unisensory noise. These findings underscore the importance of carefully dissecting the interplay between unisensory fidelity and multisensory priors in order to properly interpret the behavioral manifestations of multisensory illusions.

### 2.2 Broader Temporal Windows of Integration are not Inherently Indicative of Enhanced Integration

A widely used metric in multisensory research is the temporal window of integration (TWI), which describes the temporal interval during which inputs from distinct sensory modalities are likely to be bound into a single perceptual event. It might stand that, upon first glance, an expanded TWI in any one group or condition relative to others demonstrates heightened or advanced levels of multisensory integration. Indeed, some developmental and clinical studies have drawn precisely such a conclusion; for example, a broadening in TWIs either for children or among some clinical groups has been interpreted as pronounced multisensory binding mechanisms (e.g., Hillock, Powers, & Wallace, 2011; Hillock-Dunn & Wallace, 2012; Zhou et al., 2022).

However, a growing body of recent theoretical and empirical work indicates that broader TWIs can arise from non-integrative processes. Chief among these is reduced unisensory temporal precision: when visual or auditory timing acuity is degraded, the system may adopt a more permissive criterion for simultaneity to avoid missing genuinely coincident events (Cary et al., 2024; Horsfall et al., 2021). In this case, a large TWI reflects an adaptive response to a noisy channel, not stronger multisensory integration. A second contributor is amodal temporal priors, learned or innate expectations about event timing, which can broaden the TWI when they favor binding across larger temporal gaps (Horsfall et al., 2021; Zhu et al., 2024). For instance, Hillock et al. (2011) and Ainsworth & Bertone (2023) note that, in developmental contexts, the TWI for audiovisual stimuli decreases with age, indicating protracted refinement of temporal sensitivity across childhood and adolescence. Some earlier interpretations loosely described wider TWIs in children as akin to a developmental ‘overshoot’ or hyper-binding phase; however, this interpretation conflates TWI width with multisensory integration strength. Rather, broader TWIs are more plausibly attributable to immature unisensory temporal precision (Wang et al., 2025), and it is therefore premature to characterize children as exhibiting inherently stronger or superior multisensory integration.

A similar caution applies to clinical research: extended TWIs reported in autism spectrum disorder (ASD) and other neuropsychiatric conditions are often taken as evidence of aberrantly elevated integration (Stevenson et al., 2014; Noel et al., 2020), yet they may instead arise from reduced unisensory temporal precision or atypical temporal priors.

Taken together, these results underscore that the temporal binding window is not a thermometric readout of multisensory integration. Rather, it is a composite measure jointly and interactively, determined by (i) the reliability (*σ*) of each unisensory modality, (ii) the observer’s amodal temporal priors, and (iii) the integrative mechanism itself. Consequently, treating a broader TWI as evidence of intrinsically ‘stronger’ integration conflates integrative and non-integrative influences, such as unisensory processing limitations and domain-general expectations, and yields ambiguous inferences about the nature of multisensory perception.

To further demonstrate how unisensory precision and causal inference parameters jointly influence the temporal window of integration, we conducted a series of simulations using the 2D-BCI model proposed by Zhu et al. (2026). In total, five simulations were performed, each targeting a single parameter of interest—*p_common_*, *σ_V_*, *σ_A_*, *σ_Vt_*, or *σ_At_*—while keeping all other parameters constant. As shown in Figure 2, manipulating any one of these variables can systematically alter the shape and breadth of the TWI in a single-flash–two-beeps (1F2B) condition of the sound-induced flash illusion paradigm.

**Figure 2.**
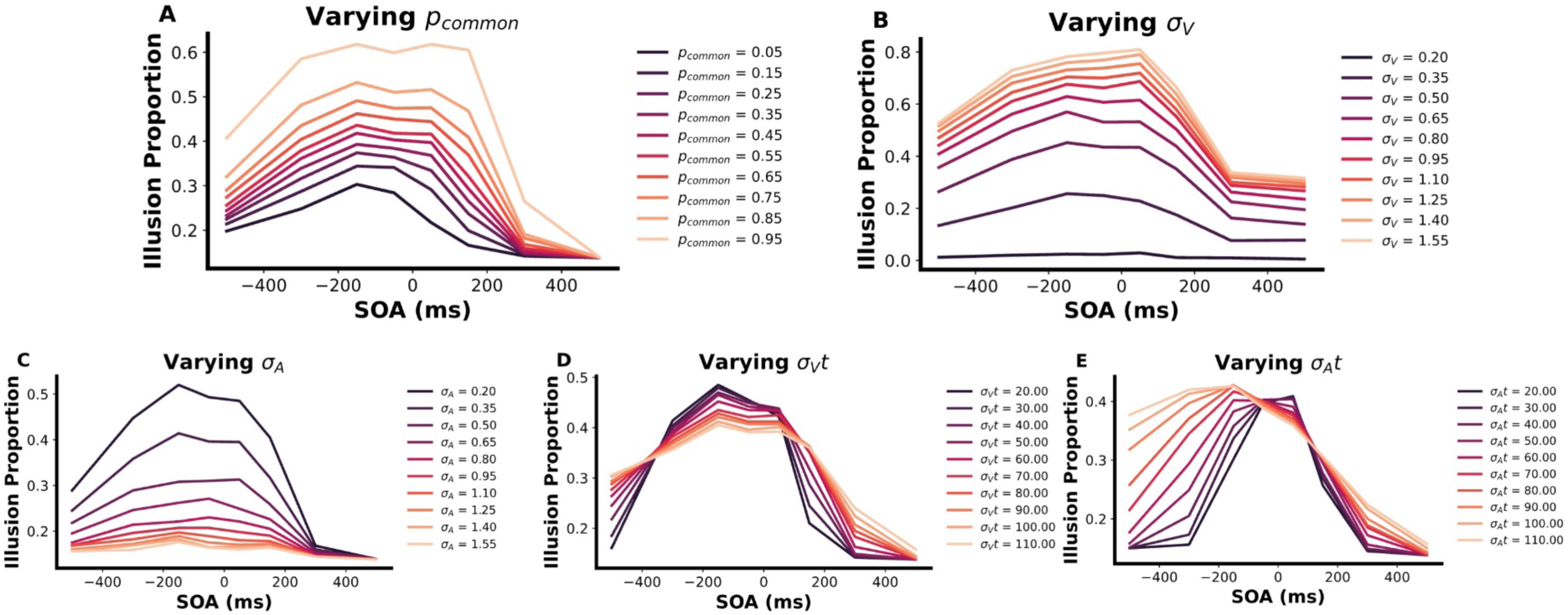
Effect of individual BCI parameters on the predicted temporal binding window for the F1B2 illusion. Simulations were performed with the two-dimensional Bayesian Causal Inference model of Zhu et al. (2024). Unless otherwise stated, parameters were fixed at *p_common_* = 0.6, *σ_V_* = 0.5, *σ_A_*= 0.3, *σ_Vt_*= 60, *σ_At_*= 40, and *μ_t_* = -50. The x-axis shows stimulus-onset asynchrony (SOA; first beep minus flash, in ms); the y-axis shows the model-predicted proportion of illusion reports (‘two flashes’) in the F1B2 condition. **(A)** Varying the causal prior *p_common_* from 0.05 to 0.95 (dark-to-light). Higher priors systematically elevate the plateau of the binding window and broaden its flanks. **(B)** Varying visual sensory noise *σ_V_* (0.20–1.55). Increasing *σ_V_* widens and raises the binding window, reflecting greater visual uncertainty. **(C)** Varying auditory sensory noise *σ_A_* (0.20–1.55). Larger *σ_A_* suppresses illusion strength and narrows the window, mirroring reduced auditory reliability. **(D)** Varying visual temporal noise *σ_Vt_* (20–110). Changes in *σ_Vt_* modulate overall amplitude but leave the window peak and width largely intact. **(E)** Varying auditory temporal noise *σ_At_* (20–110) produces a complementary pattern to panel D, again altering amplitude more than peak location or breadth. Collectively, the panels illustrate that both multisensory (*p_common_*) and unisensory (*σ* parameters) factors can reshape the temporal binding window, but their signatures differ: causal-prior manipulations primarily raise and extend the window, numerosity noise mainly produces symmetric changes in the height and width of the window without shifting its temporal center, whereas temporal-noise manipulations mainly scale its height and its center with minimal shifts in peak SOA.

Interestingly, whereas changes in *p_common_*, *σ_V_*, and *σ_A_* (i.e., the numerosity precision of each modality) tended to shift both the peak amplitude and the breadth of the TWI, variations in *σ_Vt_*, and *σ_At_* (i.e., the temporal precision of each modality) had a more selective effect. Specifically, the overall ‘height’ or amplitude of the TWI curves was sensitive to these unisensory numerosity parameters, yet the location of the peak, where the illusion proportion is highest, remained relatively stable. This finding implies that while unisensory noise can expand or contract the TWI, it does not necessarily shift the maximal point of perceived simultaneity to a large degree.

Therefore, these simulation results reinforce the notion that the TWI is not a straightforward proxy for ‘stronger’ or ‘weaker’ multisensory integration. Instead, it emerges from a complex interplay among *p_common_*, unisensory noise (both in signal strength and timing), and any amodal priors inherent to the observer. Critically, attributing changes in TWI solely to enhanced or diminished integrative capacity risks overlooking how shifts in unisensory timing precision or domain-general temporal expectations can also drive variations in apparent binding windows.

### 2.3 Different Illusory Outcomes Do Not Necessarily Reflect Distinct Integrative Mechanisms

Within certain paradigms, different sensory conditions can produce superficially contrasting illusory outcomes often described as ‘fission’ and ‘fusion.’ For instance, in auditory-visual temporal numerosity tasks, presenting multiple beeps with a single flash can sometimes lead observers to perceive more flashes than were actually shown (a ‘fission’ effect), whereas in other contexts, the same observer may underestimate the number of stimuli (a ‘fusion’ effect). At first glance, these dual manifestations may appear to reflect two separate multisensory processes: one that produces spurious splitting of events and another that merges events into a single percept.

Some researchers have independently measured fission and fusion effects within individual observers and interpreted variations in their relative magnitude across different groups or age ranges as indicative of distinct underlying multisensory mechanisms. For instance, McGovern et al. (2014) examined age-related differences in susceptibility to the sound-induced flash illusion (SiFI) and found that older adults exhibited a stronger fission illusion at longer stimulus onset asynchronies (SOAs) compared to younger adults, while fusion effects remained relatively stable across age groups. This finding was taken as evidence that separate mechanisms may govern the two illusions. Similarly, Yu et al. (2022) explored the impact of momentary rewards on the SiFI and found that reward associations selectively reduced the fission illusion without affecting the fusion illusion, leading to a similar conclusion regarding their distinct underlying processes. Additionally, Zhou et al. (2022) reported that older adults were more susceptible to the fission illusion than younger individuals, while fusion effects did not exhibit significant age-related differences.

However, these findings do not necessarily indicate that fission and fusion are mediated by distinct integrative mechanisms. Instead, a more parsimonious explanation is that both illusions emerge from a unified computational principle governed by Bayesian Causal Inference (BCI). Both behavioral and neuroimaging results demonstrate that the BCI framework posits that the brain integrates sensory information by estimating the probability that signals originate from a common or separate source, modulating perception accordingly (Rohe et al., 2019; Wozny, Beierholm, & Shams, 2010). In this view, both fission and fusion effects result from a dynamic weighting of sensory reliability and prior expectations rather than being driven by distinct neural pathways. In other words, whether an individual perceives more events (fission) or fewer events (fusion) can hinge on small shifts in sensory noise parameters or on variations in the observer’s prior beliefs about event structure, rather than on separate ‘fission pathways’ and ‘fusion pathways.’ Thus, while it can be empirically illuminating to measure fission and fusion illusions independently, interpreting any disparity between them as evidence of multiple distinct multisensory mechanisms may conflate unisensory noise or contextual priors with an entirely separate integrative architecture.

**Table 1.** Deconstructing Multisensory Constructs. This table summarizes frequent misinterpretations in multisensory integration research where observed behavioral effects are conflated with underlying neural or computational mechanisms. By identifying latent confounding variables, such as unisensory reliability and causal priors, we outline a diagnostic framework that combines experimental design with computational modeling.

| Construct | Common/In correct interpretation | How to tell apart / Required data & model (what to collect & fit) |
| --- | --- | --- |
| **Illusion magnitude**(§2.1) | ‘Bigger illusion ⇒ stronger integration.’ | **Design:** Include unisensory conditions to probe perception in each modality and to estimate basic sensory precision (e.g., $\sigma_V$ and $\sigma_A$).<br>**Model:** Fit Bayesian Causal Inference (BCI) model to estimate $p_{common}$ and sensory noise parameters separately. |
| **Temporal window of integration (TWI)**(§2.2) | ‘Wider TWI ⇒ stronger integration.’ | **Design:** Ideally, studies should include unisensory timing measures to characterize temporal precision in each modality, together with multiple bisensory conditions.<br>**Model:** Time-extended BCI (eg. Zhu et al., 2026) that separates noise, priors. |
| **Fission vs. Fusion**(§2.3) | ‘Fission and fusion reflect different mechanisms.’ | **Design:** Keep multisensory stimuli identical while shifting the observer's numerosity prior ($\mu_N$) through training or manipulation in experimental design.<br>**Model:** Fit a BCI with latent N and prior $p(N; \mu_N, \sigma_N)$; map how shifting $\mu_N$ redistributes fission and fusion.<br>**Diagnostic:** Changes in fusion versus fission can be explained by shifts in the numerosity prior alone, without requiring separate mechanisms. |

## 3 Teasing Apart Unisensory and Multisensory Factors Using Computational Models

Having discussed the ‘dont’s’ it is time to turn towards the recommended practices for multisensory psychophysical studies. An obvious recommendation is to have a plan in place for data analysis before finalizing the experimental design and submit a registered report (Chambers & Tzavellam, 2022).

A rigorous way to clarify and characterize the psychophysical data is to utilise a computational model, such as the Bayesian Causal Inference model (Figure 3A). The BCI model (Körding et al., 2007; Shams & Beierholm, 2010) assumes that participants implicitly infer the potential causal structure (e.g. one cause C=1 or two causes C=1) that gave rise to the multisensory information the nervous system receives, using Bayesian Inference:

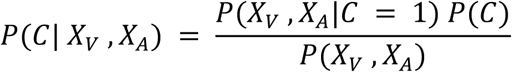

**Figure 3.**
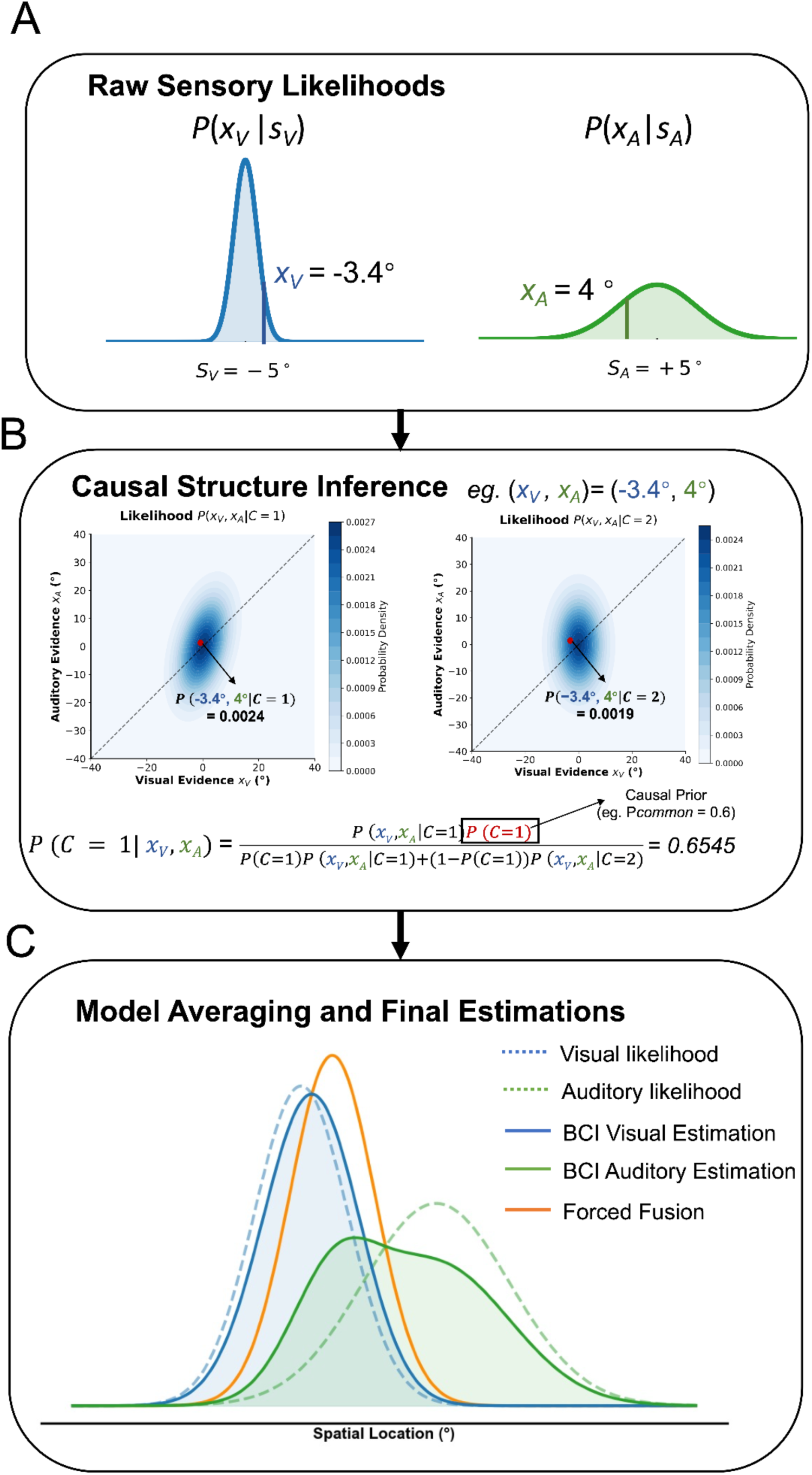
Multisensory processing computational frameworks: Bayesian causal inference. (A) Sensory Likelihoods. The system receives independent sensory signals from visual (*x_V_*) and auditory (*x_A_*) modalities, modeled as Gaussian distributions centered at their respective physical locations (*s_V_*, *s_A_*) with modality-specific sensory noise (σ_V_, σ_A_). In the illustrated example, the physical stimuli are located at *s_V_* = -5**°** and *s_A_* = +5**°**, the sensory precisions are predefined with σ_V_ **=** 3**°** (blue distribution) and σ_A_ = 10**°** (green distribution), yielding realized sensory evidence of *x_V_* **=** -3.4**°** and *x_A_***=** 4.0**°** respectively. **(B) Causal Structure Inference.** The observer evaluates the probability of a common cause (C=1) versus independent causes (C=2) for the observed sensory evidence. The observer infers the underlying causal structure by comparing the joint likelihood distributions P(*x_V_*, *x_A_*|C) under two competing hypotheses. The left heat map shows the joint distribution for a common cause (C=1), where the correlation between *x_V_* and *x_A_* is captured by a covariance matrix. The right heat map shows the joint distribution for independent causes (C=2), where the modalities are treated as orthogonal. The posterior probability of a common cause, P(C=1|*x_V_*, *x_A_*), is computed via Bayes’ rule, integrating the likelihood of the sensory evidence given both causal structures and a prior belief about common cause (*p_common_*). For the specific evidence pair (*x_V_*, *x_A_*) = (-3.4**°**, 4.0**°**), the likelihoods are computed for both hypotheses (yielding P(*x_V_*, *x_A_*|C=1) = 0.0024 and P(*x_V_*, *x_A_*|C=2) = 0.0019). Given a prior belief of *p_common_* = 0.6, the posterior probability of a common cause is calculated as P(C=1|*x_V_*, *x_A_*) = 0.6545. **(C) Model Averaging and Final Spatial Estimations.** The final spatial estimates are derived by averaging the predictions of the two causal models weighted by their respective posterior probabilities (generated by BCI toolbox (Zhu et al., 2024)). The orange curve represents the "Forced Fusion" estimate (corresponding to the assumption that all sensory signals arise from a single common source (C=1), such that cues are obligatorily integrated according to their relative reliabilities.), while the solid blue and green curves represent the BCI-derived posterior distributions for visual and auditory spatial estimates, respectively. Dashed lines denote the sensory likelihoods for reference.

Based on this inferred causal structure, perceptual estimates are computed based onthe probability of the alternative causal hypotheses. Final behavioral reports can be generated via different decision strategies, such as model averaging, model selection, or probability matching (Wozny et al., 2010), each yielding distinct but formally defined predictions over response distributions.

Given a participant’s response matrix, the free parameters Θ={*p_common_*,*σ_A_,σ_V_*,*σ_prior_*} can be estimated via maximum-likelihood optimization or more advanced machine-learning routines such as Bayesian Adaptive Direct Search (BADS; Acerbi & Ma, 2017) and Variational Bayesian Monte Carlo (VBMC; Acerbi, 2020). More complex experimental settings involving multiple stimulus dimensions (e.g., space, time, number) can be addressed using multidimensional extensions of the BCI framework (Zhu et al., 2026).

One of the biggest advantages of using a model such as BCI is that it teases apart the contributions of the different computational factors such as prior expectations, versus unisensory precisions, versus the binding/integration across the senses, the decision-making strategy, etc.. Each of these factors reflect part of the neural process assumed to have generated the recorded data. As we have highlighted above, without a computational model it can be very difficult to identify the factor underlying the observed differences in behaviour across individuals, populations, or experimental settings. One would not generally be able to distinguish between a multisensory factor (such as *p_common_*), unisensory factors (such as the likelihood variances) or amodal factors (such as an amodal spatial prior variance). Recent modelling work has begun to explicitly decompose multisensory behaviour into underlying components, demonstrating, for instance, how developmental changes in illusion susceptibility can be accounted for by shifts in unisensory reliability (Wang et al., 2025), and how clinical differences can be dissociated at the level of distinct computational parameters (Cao et al., 2019). Similar computational frameworks have also been used to link neural mechanisms to temporal integration processes, demonstrating how physiological factors can selectively modulate temporal binding without necessarily altering other components of the inference process (e.g., D’Angelo et al., 2026).

**BOX 1** | Practical Guidelines for Applying Bayesian Causal Inference (BCI) in Multisensory Research

While fitting the BCI to data has previously required the ability to program (in Matlab, Python etc.), and possibly writing novel code, a recently developed toolbox can be used for beginners with little to no programming experience (Zhu et al., 2024). However, to use BCI properly, the experimental design needs to accommodate proper parameter estimation:

- Include both unisensory and multisensory blocks: ideally the experiment will consist of both unisensory and multisensory conditions. This allows for the model to be fitted across a variety of sensory conditions, allowing for more robust parameter fitting. Unisensory trials anchor the sensory-noise parameters (σA, σV) and constrain amodal priors; multisensory trials constrain the causal prior *P*common and amodal priors. Practical tips: Interleave or counter-balance A-only, V-only, and A + V conditions within each session. Interleaving the trials would reduce the likelihood of distinct response criteria across sensory conditions.
- Have subjects report both modalities on each trial: In experimental conditions where both modalities/senses provide useful responses it can be useful to have subjects report both responses within the same trials. This provides data on joint probabilities of the two sensory modalities, allowing a more reliable parameter estimation. However, in situations where one modality is much stronger than the other, and at performance ceiling, collecting data on that modality would not be necessary for improving parameter estimation. Practical tip: Omit the recording of the stronger modality if pilot data show ceiling-level accuracy; otherwise record both.
- Large enough #trials/condition: As with fitting any parameters to a model (whether a simple t-test or a complicated hierarchical Bayesian model), sufficient number of trials is important for reliable parameter fitting. Hard guidelines are difficult to provide, but we find a minimum of 80 total trials per free parameter (with a minimum of about 15 trials per sensory condition) is useful when working with the BCI model. For the four-parameter BCI core (σA, σV, σprior, *p_common_*), aim for ≥ 300 analysable trials per participant, distributed across discrepancy levels (e.g., a minimum of 15 trials/condition x 24 conditions).

## 4 Alternative Frameworks for Multisensory Processing

### 4.1 Multisensory Correlation Detection

Whereas BCI serves as a normative computational model that formalizes how latent causal structures and priors should be integrated, the Multisensory Correlation Detector (MCD) (Parise & Ernst, 2016) offers a stimulus-computable process model of how such inferences might be algorithmically implemented. In the MCD, audiovisual inputs pass through modality-specific temporal filters and linear combinations to yield two time-resolved outputs: a correlation signal (how strongly the streams co-vary) and a lag signal (which stream leads). With only a few temporal constants, the MCD can predict synchrony/temporal-order and causality judgments, and it reproduces observers’ classification images with high fidelity, suggesting that correlation detection may constitute an elementary algorithmic step that feeds higher-level decisions about whether cues share a common cause. Importantly, recent MEG work shows that brain dynamics track MCD components (Pesnot Lerousseau et al., 2022), supporting its neurobiological plausibility. In practice, these models operate at different levels of explanation: MCD provides a mechanistic, dynamic implementation (Parise & Ernst, 2025), whereas BCI provides a normative computational account that can characterize priors, unisensory noise, and binding tendencies.

### 4.2 Drift Diffusion Models

When both reaction time and accuracy are available, and a speed-accuracy trade-off exists in the behavioral data, drift diffusion models (DDM) can be used to account for the behavior (Ratcliff & Rouder, 1998). DDMs are a family of models, some of which are normative, based on the premise that the nervous system integrates information over time before reaching a decision. There are existing toolboxes for fitting DDMs to experimental data, simplifying their use greatly (Wiecki et al., 2013).

The simplest version of the DDM assumes that a subject has performed a binary choice (e.g. ‘did you hear a beep’ after being exposed to auditory noise with a potential beep embedded in it) and that choice and reaction time has been recorded. The decision making of the subject is framed as an evidence accumulation process, by which the subject keeps gathering stochastic information for either option (‘yes’ or ‘no’) until an information threshold is reached. The time to reach this threshold (plus a potential non-decision time) constitutes the reaction time of a trial. After gathering responses and reaction times across a large number of trials it is possible to fit parameters of this model to the data. The two main parameters are the drift diffusion rate (how fast information accumulates) and the threshold, although extensions of the model have introduced other variables (e.g. non-decision time, or biased starting point). By fitting parameters for each experimental condition, it is possible to test the change in a given parameter across sensory conditions. For example, an experimenter can use the DDM model to ask whether an added stimulus (e.g. visual flash) synchronised with the auditory stimulus causes the subject to accumulate information faster (see Drugowitsch et al., 2014; Franzen et al., 2020; Chau et al., 2021 as examples). Without such computational models, it can be difficult to disentangle different contributions to reaction time, such as the effect of improved discriminability versus increased bias. Indeed, studies have shown DDM can reveal benefits of multisensory processing even when accuracy data or reaction time data alone would not (Drugowitsch et al., 2014; Franzen et al., 2020; Chau et al., 2021; Murray et al., 2020). For more complicated tasks it may be necessary to develop bespoke models using more advanced statistical tools (e.g. Stan, for an example see Henrich et al., 2023). Importantly, these models do not tease apart unisensory reliabilities and tendency to integrate the senses as underlying factors of interaction between the two modalities.

### 4.3 Statistical facilitation accounts of multisensory speed-up

Alongside DDMs, a large body of multisensory reaction time studies use redundant-signals tasks, where responses are faster to bisensory than unisensory cues (the redundant-signals effect, RSE; Otto & Mamassian, 2017). Two canonical accounts explain this facilitation. Race model (aka, separate activation models) attributes the RSE to statistical facilitation: whichever unisensory channel finishes first triggers the response (Raab, 1962; Miller, 1982). However, neural–behavioural evidence suggests that audiovisual detection may instead involve partially independent evidence accumulators whose outputs jointly activate a downstream motor decision stage, providing a mechanistic account of multisensory speed-up beyond simple probability summation (Egan et al., 2025). A standard diagnostic is Miller’s race-model inequality (RMI), which compares the empirical bimodal RT distribution with the summed unisensory distributions; violations of the RMI imply that probability summation is insufficient (Miller, 1982; Gondan & Minakata, 2016). Beyond the classic RMI logic, Diederich and colleagues developed co-activation formalisms that go beyond probability summation. In a series of papers, Diederich (1995) quantitatively evaluated counter and diffusion co-activation models, and with Colonius introduced the Time-Window-of-Integration (TWIN) model, which links super-additive RT facilitation to a finite temporal window within which cross-modal evidence is pooled (Diederich & Colonius, 2004; Diederich & Colonius, 2015). This approach provides a mechanistic model when RMI violations indicate that a race architecture is insufficient to account for the reaction time data. Recent work generalizes these tools (e.g., testing the RMI in difficult/error-prone tasks or with more than two modalities; Colonius, 2017; Gondan, Dupont, & Blurton, 2020) and shows cases where race models suffice for audiovisual motion decisions (Chua et al., 2022).

These studies have greatly advanced the study of reaction times in multisensory integration and provided robust ways to quantify multisensory gains. However, there is still no unified, cross-task computational framework that can reliably detect and quantify the degree of integration when analyzing RT data. More importantly, these models do not tease apart unisensory reliabilities and tendency to integrate the senses as underlying factors of interaction between the two modalities.

## 5. Discussion

Multisensory integration research is at a turning point. For decades, the field has relied on heuristic measures, for example, crossmodal illusion magnitude, width of the temporal binding window as proxies for ‘integration strength.’ These measures have been invaluable for charting the phenomenology of MSP, but they are systematically misinterpreted when considered in isolation. Going forward, MSP research should follow model-based inference pipelines rather than descriptive heuristics and explicitly separate unisensory fidelity, amodal priors, and integrative mechanisms. This would offer a more principled framework for interpreting group differences across development, clinical, and aging populations and link the most basic cognitive science with translation domains, from neuropsychiatric diagnostics to building robust sensor fusion in AI and robotics.

In this review, we have highlighted several common pitfalls in the interpretation of multisensory processing data, including the reliance on illusion magnitude, temporal binding window width, or performance facilitation as direct proxies for ‘integration strength.’ Through targeted simulations, we demonstrated how these descriptive measures can arise from distinct underlying factors, such as changes in unisensory reliability, or shifts in amodal priors, without necessarily reflecting a genuine change in integrative tendency. Within this context, Bayesian Causal Inference (BCI) (Körding et al., 2007; Shams & Beierholm, 2010; 2022) provides a particularly useful general framework. By explicitly modeling latent causal structure and formally separating sensory likelihoods from causal priors, BCI enables a principled decomposition of observed behavior into interpretable underlying factors (Körding et al., 2007; Shams & Beierholm, 2010). This factorization allows researchers to distinguish whether an observed behavioral change reflects altered unisensory fidelity, shifts in amodal expectations, or a genuine difference in binding propensity. Crucially, this separation is what permits rigorous comparison across developmental stages, clinical populations, or experimental manipulations without conflating distinct sources of variability.

Beyond BCI, several influential computational frameworks have substantially advanced the field. Sequential-sampling models such as the Drift Diffusion Model (DDM) have provided a powerful account of how multisensory information enhances decisions by modulating evidence accumulation dynamics (Ratcliff & McKoon, 2008; Drugowitsch et al., 2014; Chau et al., 2021; Murray et al., 2020). Empirically, multisensory conditions often exhibit increased drift rates or reduced non-decision times, consistent with facilitation of evidence processing. Likewise, stimulus-computable accounts such as the Multisensory Correlation Detector (MCD) (Parise & Ernst, 2016; Pesnot Lerousseau et al., 2022), which operate directly on raw sensory inputs rather than abstract variables, have offered mechanistic insight into how temporal correlation signals may feed higher-level judgments of common cause. At the behavioral-performance level, statistical facilitation models, including Race Model Inequality and Co-activation accounts, have rigorously formalized redundancy gains and provided conservative benchmarks for multisensory speed-up effects (Raab, 1962; Miller, 1982). These models have been instrumental in moving the field beyond purely descriptive phenomenology and toward quantitative explanations. However, despite their explanatory power, these frameworks do not explicitly decompose multisensory interaction into separable causal components. For example, in DDM-based analyses, an increased drift rate under multisensory stimulation clearly reflects enhanced information processing. But if one individual exhibits a bigger increase in drift rate in multisensory vs unisensory conditions compared to another individual, the stronger multisensory benefit in drift rate can stem from different factors. Such disparity does not specify whether the effect arises from a difference in prior tendency to integrate (analogous to a difference in *Pcommon*), or from a difference in relative unisensory uncertainties. Observing an elevated drift rate in the multisensory condition is analogous to observing a robust multisensory illusion: it confirms interaction but does not clarify its source. In other words, the model quantifies how much performance changes, but not why the system favors integration in a given context. The same structural limitation applies to MCD and Race/Co-activation frameworks as well.

While BCI provides a principled framework for decomposing multisensory behavior, important open questions remain. Current frameworks are largely tailored to choice or perceptual reports and often abstract away from reaction-time dynamics, leaving open the question of how causal inference unfolds over time within evidence accumulation processes. Integrating BCI with sequential-sampling frameworks could provide a principled way to link latent causal structure to decision dynamics. In addition, multisensory perception does not occur in isolation: attention, working memory, learning, and task context all shape how sensory information is weighted and bound. Future work should therefore investigate how these cognitive factors modulate causal priors and sensory likelihoods within a unified inferential framework. Extending BCI to account for temporal dynamics, higher-order cognitive influences, and complex naturalistic stimuli will be essential for capturing the richness of real-world perception. By situating causal inference within a broader cognitive architecture, multisensory research can move toward models that are not only explanatory but also integrative across levels of analysis (Shams & Beierholm, 2022). Such alignment will foster cumulative progress, reduce interpretational ambiguity, and enable more coherent dialogue between basic research, clinical investigation, and artificial systems design.

## Glossary

| Symbol | Term | Description |
| --- | --- | --- |
| $x_A, x_V$ | Sensory Measurements | Noisy sensory measurements of auditory and visual events. |
| $\sigma_V, \sigma_A$ | Sensory Noise | Standard deviations of auditory and visual sensory noise. |
| $\sigma_{Vt}, \sigma_{At}$ | Sensory Noise (Temporal) | Standard deviations of auditory and visual temporal noise. |
| $\mu_p, \sigma_p$ | Numerosity Prior | Mean and standard deviation of the numerosity prior. Here illustrated using a temporal numerosity judgment task, but the parameterization generalizes to priors over other stimulus dimensions (e.g., spatial location). |
| $\mu_{tp}, \sigma_{tp}$ | Temporal Prior | Mean and standard deviation of the temporal prior (shared across modalities). |
| C | Causal Structure | Latent causal variable (1 = common cause; 2 = independent causes). |
| $p_{common}$ | Causal Prior | Causal prior: probability that the two signals arise from a common cause. |
| SOA | Stimulus Onset Asynchrony | The temporal delay between the presentation of visual and auditory stimuli. |
| Model Averaging | - | An integration strategy where estimates from both causal structures are weighted by their respective posterior probabilities. |
| TBW | Temporal Binding Window | The time interval within which different sensory inputs are likely to be integrated as a single event. |

